# Potentiation of Adipogenesis by Reactive Oxygen Species is a Unifying Mechanism in the Pro-adipogenic Properties of Bisphenol A and its New Structural Analogues

**DOI:** 10.1101/2022.09.08.507176

**Authors:** Radha D. Singh, Jessica L. Wager, Taylor B Scheidl, Liam T. Connors, Sarah Easson, Mikyla A. Callaghan, Samuel Alatorre-Hinojosa, Lucy H. Swift, Pina Colarusso, Anshul Jadli, Timothy E. Shutt, Vaibhav Patel, Jennifer A. Thompson

## Abstract

**Aims:** Structural analogues of bisphenol A (BPA), including BPS and BPF, are emerging environmental toxicants as their presence in the environment is rising since new regulatory restrictions were placed on BPA-containing infant products. The adipogenesis-enhancing effect of bisphenols may explain the link between human exposure and metabolic disease; however, underlying molecular pathways remain unresolved.

**Results:** Exposure to BPS, BPF, BPA or ROS generators enhanced lipid droplet formation and expression of adipogenic markers after induction of differentiation in adipose-derived progenitors isolated from mice. RNAseq analysis in BPS-exposed progenitors revealed modulation in pathways regulating adipogenesis and responses to oxidative stress. ROS was higher in bisphenol-exposed cells, while co-treatment with antioxidants attenuated adipogenesis and abolished the effect of BPS. There was a loss of mitochondria membrane potential in BPS-exposed cells and mitochondria-derived ROS contributed to potentiation of adipogenesis by BPS and its analogues. Male mice exposed to BPS during gestation had higher whole-body adiposity, as measured by TD-NMR, while postnatal exposure had no impact on adiposity in either sex.

**Innovation:** These findings support existing evidence showing a role for ROS in regulating adipocyte differentiation and are the first to highlight ROS as a unifying mechanism that explains the pro-adipogenic properties of BPA and its structural analogues.

**Conclusion:** ROS act as signaling molecules in the regulation of adipocyte differentiation and mediate bisphenol-induced potentiation of adipogenesis.

## INTRODUCTION

The *Lancet* Commission on pollution and health identified chemical intensification of the environment as one of the most significant causes of premature death (Landrigan et al., 2018). The plastic industry is a major contributor to the environmental chemical burden, emitting 400 million tonnes of synthetic chemicals each year (Naidu et al., 2021). Bisphenols, plasticizers used in the manufacturing of polycarbonate plastics and epoxy resins, are classified as endocrine disrupting chemicals (EDC), which interfere with developmental and reproductive processes by mimicking endogenous ligands to receptors and acting as agonists or antagonists. After regulatory bans on the sale or import of baby products containing bisphenol A (BPA), manufacturers responded by replacing it with structural analogues, marketing them as ‘safe’ substitutes. However, evidence supporting this claim of health benefit is lacking and emerging data suggest that BPA analogues are another example of ‘regrettable substitution’, where toxic chemicals are replaced with equally toxic substitutes (Zimmerman and Anastas, 2015). BPA substitutes, such as bisphenol S (BPS) and bisphenol F (BPF), are now recognized as emerging toxicants as their production is on the rise, approaching or exceeding the world average daily intake of BPA in countries that introduced regulatory restrictions (Wang et al., 2020). Therefore, there is a need to fill the gap in knowledge with respect to health effects and biological properties of the new BPA substitutes.

A large body of published data show that BPA promotes adipogenesis *in vitro* (Ariemma et al., 2016, Ohlstein et al., 2014). Fewer studies have investigated the structural analogues of BPA; however, the available evidence to-date reveal that commonly used substitutes, BPS and BPF, exhibit similar pro-adipogenic properties as their predecessor, with some studies showing effects at lower doses (Ramskov Tetzlaff et al., 2020). Investigation into the molecular pathways mediating the adipogenic effect of bisphenols have almost exclusively focused on endocrine pathways as it is well known that BPA and its analogues bind to estrogen, androgen and other hormone receptors. These studies have yielded inconsistent results with some showing a role for estrogen receptors and others a role for glucocorticoid receptor (Ahmed and Atlas, 2016, Boucher et al., 2016). Overall, the cellular mechanisms responsible for the adipogenesis-promoting properties of bisphenols remain unresolved.

Adipogenesis is a sequential differentiation process by which mesenchymal stem cells commit to the adipocyte lineage and terminally differentiate into lipid-storing adipocytes. Specification of stem cells to the adipocyte fate in subcutaneous adipose tissue (SAT), the primary site of energy storage, is restricted to late fetal life (Jiang et al., 2014, Wang et al., 2013). Establishment of the progenitor pool is followed by two periods of rapid fat accumulation occurring after birth and during puberty, with adipocyte numbers stabilizing thereafter (Holtrup et al., 2017). In adult adipose depots, *de novo* adipogenesis is stimulated to support normal cellular turnover and to increase lipid storage capacity in response to an obesogenic environment (Jeffery et al., 2015, Wang et al., 2013, White and Ravussin, 2019). Current evidence suggests that new adipocytes arising in adult depots derive from a distinct compartment of progenitors that are also specified to the adipocyte lineage *in utero* (Jiang et al., 2014, Wang et al., 2015). Thus, the early life window of adipogenesis establishes the setpoint of adiposity and programs lipid buffering capacity in adult depots.

Herein, we set out to determine the effect of BPS on adipogenesis and elucidate the molecular pathways involved. Our data reveal a critical role for reactive oxygen species (ROS) in acting as physiological signaling molecules in the regulation of adipogenesis under normal conditions. Increased ROS production, contributed by mitochondria dysfunction, was responsible for the potentiation of adipogenesis in progenitors exposed to BPS as well as other structural analogues. Thus, bisphenol-induced ROS may be a unifying mechanism that explains the pro-adipogenic properties of BPA and its substitutes. Second, we exposed mice to an environmentally relevant dose of BPS during or after the early life window of adipogenesis and determined that only early-life exposure had an impact on adiposity in adulthood. Therefore, exposure to BPS before birth may raise the setpoint for adiposity, predisposing to obesity.

## RESULTS

### BPS, BPF and BPA potentiate differentiation of adipocyte progenitors *in vitro*

Adipocyte progenitors isolated from inguinal SAT (iSAT) were exposed to various concentrations of BPS prior to differentiation. Exposure of progenitors to BPS increased lipid droplet formation on day 2, 4 & 7 post-differentiation, albeit in a non-monotonic manner such that the increase in differentiation at low doses was attenuated at the highest dose (Figure 2A & B). The MTT assay revealed a decrease in cell viability at 25 μM, the highest dose of BPS (Supplementary Figure 1), suggesting that the attenuation of lipid droplet staining at this concentration of BPS is due to cytotoxicity. On day 7 of differentiation, lipid droplet staining was higher in progenitors exposed to BPA or BPF (Supplementary Figure 2A &B). The adipogenic response to BPA occurred at higher doses, while the attenuation of the adipogenic response at 25 μM of BPS was not observed at the same concentration of BPA or BPF. On day 2 of differentiation in BPS-exposed progenitors, there was an increase in mRNA expression of peroxisome proliferator-activated receptor gamma (*Pparγ*), a nuclear receptor required for adipogenesis, as well as glucose transporter type 4 (*Glut4*), an adipogenic marker (Figure 2C). On day 2, the mRNA expression of CCAAT/enhancer binding protein beta (*C/ebpβ*), an early regulator of differentiation, was decreased in BPS-exposed progenitors (Figure 2C). *Glut4* mRNA expression remained high on day 4 and there was an increase in stearoyl-CoA desaturase 1 (*Scd1*), a key enzyme involved in lipogenesis (Figure 2C). Protein expression of adiponectin, SCD1, fatty acid binding protein 4 (FABP4) and C/EBPβ was increased on day 2 and 4 of differentiation in BPS-exposed progenitors (Figure 2D & E).

**Figure 1:**
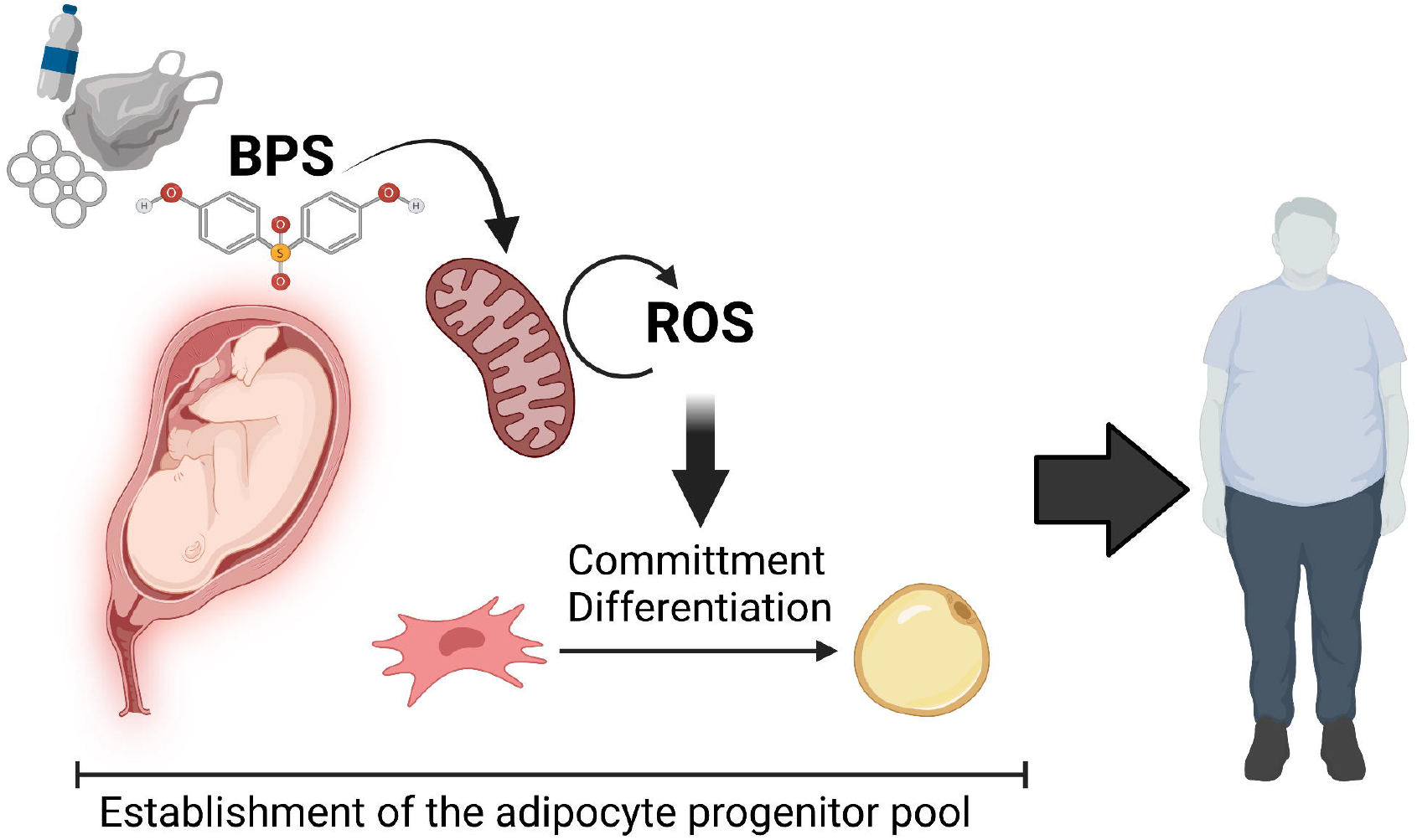
Summary Graphic Illustration. Reactive oxygen species (ROS) act as signaling molecules that regulate the commitment and differentiation of adipocyte progenitors, a developmental event that occurs during a discrete window in intrauterine life and establishes the setpoint of adiposity. Bisphenol S (BPS) potentiates adipogenesis via stimulating an increased production of ROS in part through inducing mitochondria dysfunction. Therefore, gestational exposure to BPS and other structural analogues of BPA may predispose to the development of later-life obesity.

**Figure 2:**
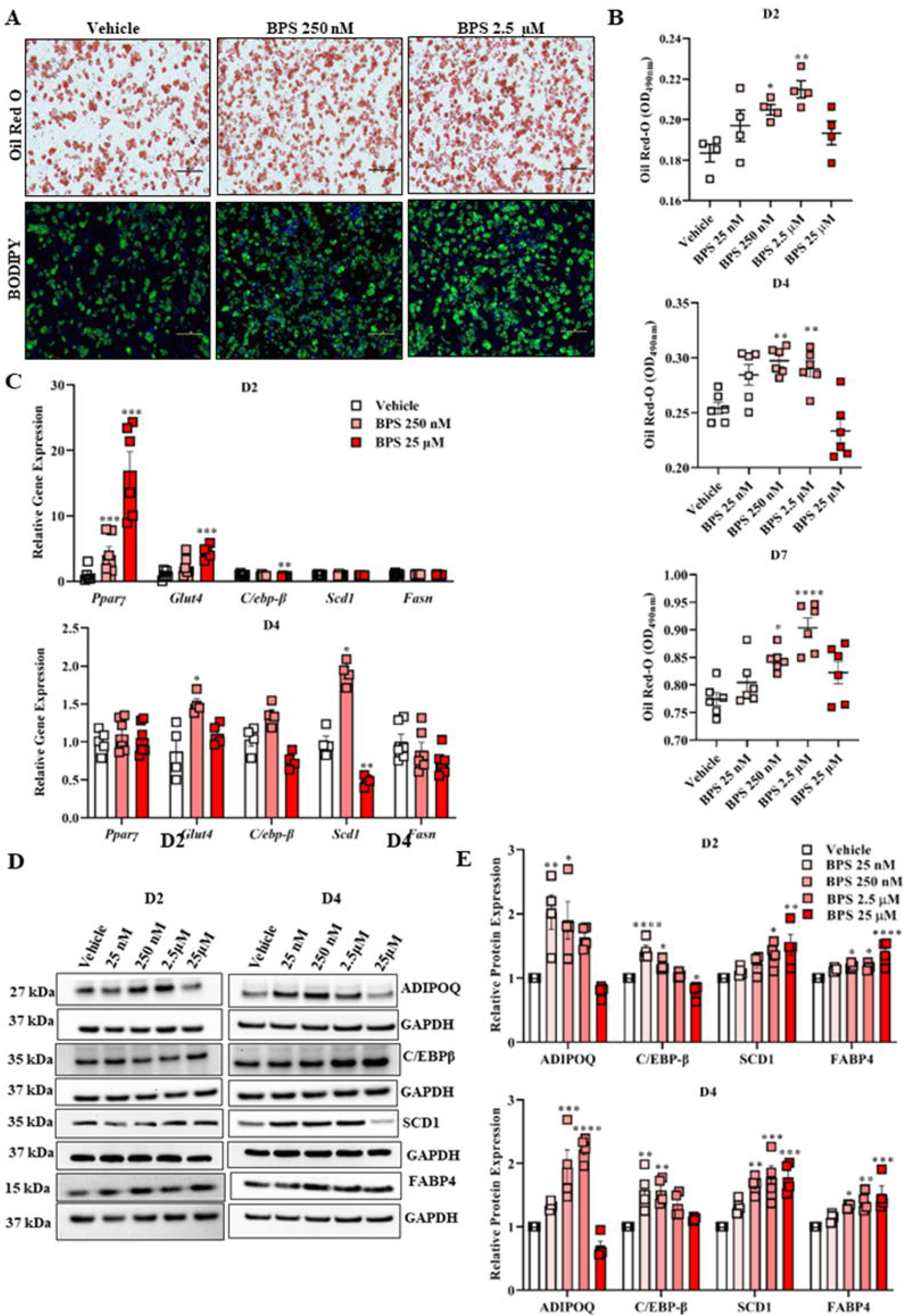
BPS potentiates differentiation of adipocyte progenitors. Representative images of lipid droplet staining by Oil Red-O or BODIPY/DAPI on day 7 of differentiation in adipose-derived progenitors exposed to BPS or vehicle control (A). Lipid droplet formation quantified by measuring the optical density (OD, 490 nm) of eluted Oil Red-O dye from stained progenitors on day 2, 4 & 7 of differentiation after BPS exposure (B). Relative mRNA expression of adipogenic markers on day 2 & 4 of differentiation (C). Representative blots (D) and quantification (E) by densitometry of Western blots for determination of protein expression of adipogenic markers on day 2 & 4 of differentiation. Each experiment was conduced in cells isolated from 2-3 animals and triplicate absorbance readings from each well are averaged. Differences are compared by One-way ANOVA with Dunnett’s multiple comparison test; * p < 0.05 BPS vs. vehicle. Abbreviations: Adiponectin (ADIPOQ); CCAAT/enhancer binding protein beta (C/EBPβ); fatty acid binding protein 4 (FABP4); fatty acid synthase (FASN); glucose transporter type 4 (GLUT4); peroxisome proliferator-activated receptor gamma (PPARγ); stearoyl-CoA desaturase 1 (SCD1).

### BPS exposure modulates gene pathways involved in oxidative stress responses

RNAseq analysis was performed in extracted RNA collected from adipose-derived progenitors on D0, at which time changes in gene expression patterns and epigenetic signatures stimulated by contact inhibition prime the cells for differentiation (Guo et al., 2016). There were over 30 DEGs involved in adipogenesis and over 12 genes involved in Nrf-2-mediated responses to oxidative stress between vehicle-treated and BPS-exposed cells (Figure 3A). MSigDB Hallmark Pathway Analysis showed that DEGs were enriched in pathways regulating cellular stress (reactive oxygen species; hypoxia), inflammatory/immune cascades (TNFα/NF-κB; IL6/JAK/STAT3; Interferon α/γ), responses to DNA damage (UV light response; G2-M checkpoint), as well as cell growth and death (p53 pathway; KRAS signaling; apoptosis; mTORC1) (Figure 3B). Interaction Networks Analysis shows the predicted relationship between DEGs involved in pathways regulating adipogenesis and Nrf2-mediated oxidative stress responses (Figure 3C). Validation by qPCR of DEGs identified by RNAseq with fold changes >1 confirmed a downregulation of antioxidant genes, catalase (*Cat*) and superoxide dismutase 3 (*Sod3*), and upregulation of *c-FOS*, a redox-sensitive early response gene involved in the regulation of cellular proliferation and differentiation (Figure 3D).

**Figure 3:**
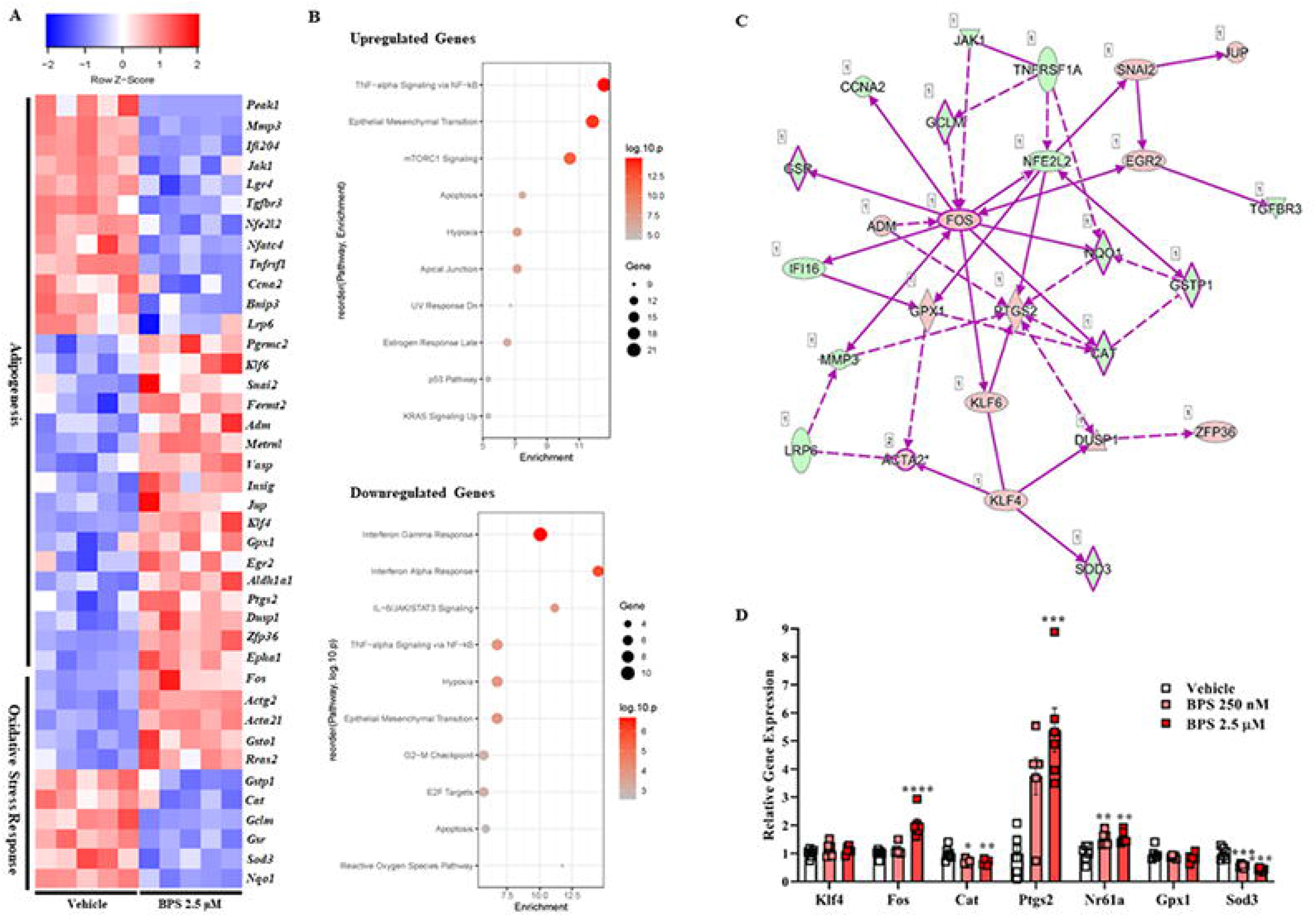
Prior to differentiation, pathways involved in adipocyte differentiation and oxidative stress are modulated in adipose-derived stem cells exposed to BPS. Heat map showing DEGs involved in adipogenesis and Nrf2-mediated responses to oxidative stress, in vehicle vs. BPS-exposed undifferentiated progenitors (A). Bubble plot showing number of upregulated and downregulated genes in the top 10 enriched pathways in MSigDB Hallmark analysis (B). IPA showing integration of DEGs involved in pathways regulating adipogenesis and NRF2-mediated responses to oxidative stress (C); Green = downregulated genes; Orange = upregulated genes. PCR validation of differences in gene expression with > 1fold change between vehicle and BPS-exposed progenitors (D). RNAseq analysis included n = 5 samples/group. * p < 0.05 BPS vs. vehicle. Abbreviations: Catalase (Cat); Kruppel-like factor 4 (Klf4); AP-1 transcription factor subunit (Fos); Glutathione peroxidase 1 (Gpx1); nuclear receptor sub-family 6 group A member 1 (Nr6a1); prostaglandin endoperoxidase synthase 2 (Ptgs2); superoxide dismutase 3 (Sod3).

### ROS production is increased in progenitors exposed to bisphenol analogues

In undifferentiated progenitors a surge in ROS generation was observed in response to acute BPS exposure in a concentration-dependent manner, as assessed by quantification of DCFDA (Figure 4A). Similarly, detection of DCFDA by fluorescence spectroscopy revealed increased ROS production after acute exposure to BPA (Figure 4B) or BPF (Figure 4C). The number of cells stained with DCFDA was quantified by flow cytometry, also showing increased ROS production after acute BPS exposure (Figure 4D & E). After 48 hrs of BPS exposure, staining intensity of CellROX™ deep red, a fluorescent probe that detects oxidative stress, was higher in BPS-exposed cells (Figure 4F & G). Similarly, CellROX staining was increased in undifferentiated progenitors exposed to BPA (Figure 4H) or BPF (Figure 4I) for 48 hrs. These data show that BPA and its structural analogues stimulate an increase ROS production.

**Figure 4:**
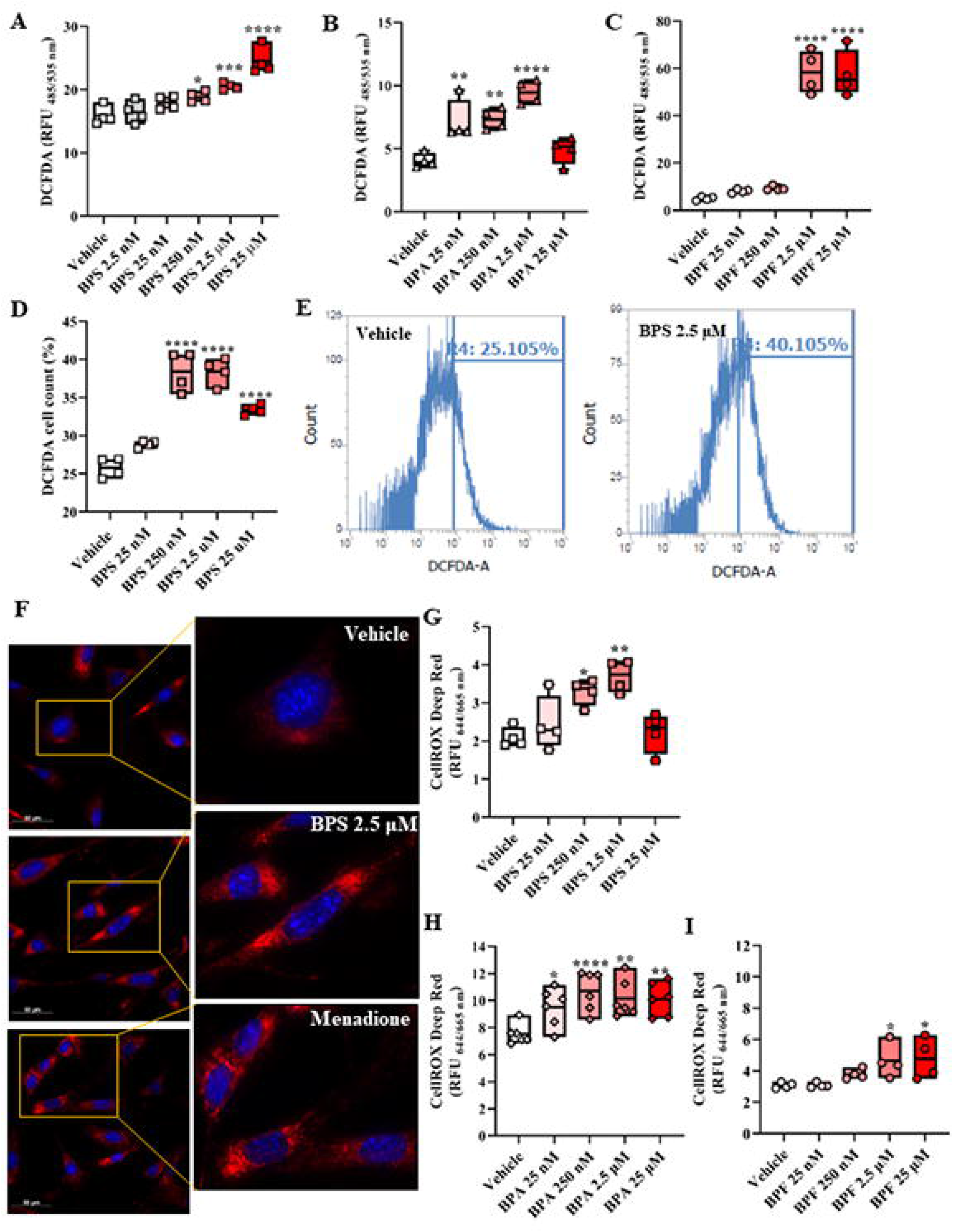
BPS and other bisphenol analogues increase production of ROS. Production of superoxide (O_2_^−^) was detected by measuring relative fluorescence units (RFU) of 2′,7′-dichlorodihydrofluorescein diacetate (DCFDA) staining by the plate reader method in undifferentiated cells acutely exposed to various concentrations of BPS (A), BPA (B) or BPF (C). Additionally, DCFDA staining was quantified by flow cytometry in progenitors exposed to BPS (D) and representative histograms are shown (E). Oxidative stress was detected in cells exposed to BPS for 48 hrs by CellROX staining. Representative images of CellROX-stained cells with DAPI nuclear staining are shown (F), and CellROX fluorescence was quantified by spectroscopy (G). Oxidative stress measured by quantification of CellROX staining was also detected in stem cells exposed to BPA (H) or BPF (I). Each experiment was conducted in cells isolated from 2-3 animals and triplicate absorbance readings from each well are averaged. Differences are compared by One-way ANOVA with Dunnett’s multiple comparison test. * p < 0.05 BPS vs. vehicle.

### ROS regulate adipogenesis and mediate BPS-induced potentiation of adipogenesis

Exposure of progenitors to low concentrations of the ROS generator, menadione, augmented lipid droplet formation on day 4 post-differentiation in a non-monotonic manner similar to BPS, such that increased lipid droplets at lower concentrations was attenuated with higher concentrations (Figure 5A). On day 2 post-differentiation in progenitors exposed to menadione, mRNA expression of *Pparγ* and *Glut4* was increased, while expression of *C/ebpβ* was decreased (Figure 5B). Similarly, exposure of progenitors to another ROS generator, 2,3-dimethoxy-1,4-napthalenedione (DMNQ), lead to a non-monotonic increase in lipid droplet staining on day 4 (Figure 5C) and an increase in mRNA expression of *Pparγ* and *Glut4* on day 2 (Figure 5D). In the absence of BPS, suppression of ROS generation via a ROS scavenger (Tempol), led to a concentration-dependent decrease in lipid droplet staining after differentiation (Figure 5E). Co-treatment of progenitors with either Tempol or the antioxidant, apocynin, abolished the effect of BPS on adipogenesis (Figure 5F-H). Together, these data reveal ROS to be critical regulators of adipogenesis and mediators of BPS-induced potentiation of adipogenesis.

**Figure 5:**
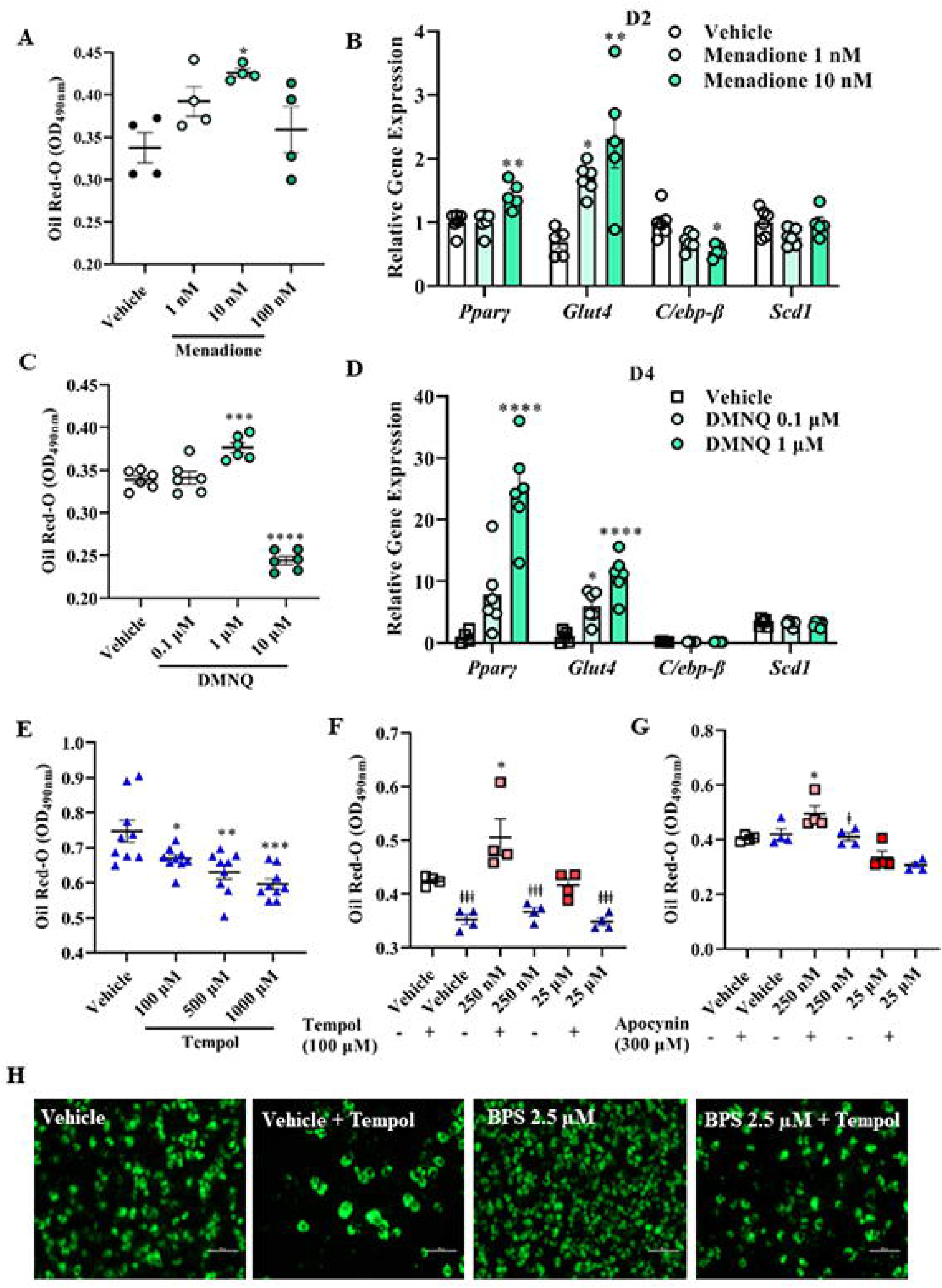
ROS regulate adipogenesis and mediate BPS-induced potentiation of adipogenesis. Lipid droplet formation assessed by measuring optical density (OD, 490 nm) of eluted Oil Red-O dye in progenitors exposed to the superoxide (O_2_^−^) generator, menadione, on day 4 of differentiation (A). Relative mRNA expression of adipogenic markers on day 2 of differentiation after exposure to menadione (B). Quantification of lipid droplet staining by Oil Red-O on day 4 of differentiation of progenitors treated with the O_2-_ generator, 2,3-dimethoxy-1,4-napthalenedione (DMNQ) (C). Relative mRNA expression of adipogenic markers on day 2 of differentiation after exposure to DMNQ (D). Quantification of Oil Red-O staining in progenitors exposed to various doses of the ROS scavenger, Tempol, in the absence of BPS (E). Absorbance of Oil Red-O staining in vehicle or BPS-exposed progenitors in the presence or absence of Tempol (F) or the NADPH oxidase inhibitor and antioxidant, apocynin (G). Representative BODIPY-stained differentiated progenitors exposed to vehicle or BPS in the presence or absence of Tempol (H). Each experiment was conducted in cells isolated form 2-3 animals and triplicate absorbance readings from each well were averaged. Differences were assessed by One-way ANOVA with Dunnett’s multiple comparison test to compared BPS vs. vehicle (* p < 0.05) or One-way ANOVA with Tukey’s post hoc test to compare vehicle or BPS with inhibitor vs. without inhibitor (‡ p < 0.05 vs. without inhibitor). Abbreviations: CCAAT/enhancer binding protein beta (C/ebpβ); glucose transporter type 4 (Glut4); peroxisome proliferator-activated receptor gamma (Pparγ); stearoyl-CoA desaturase 1 (Scd1).

### Dysfunctional mitochondria are important sources of ROS in BPS-exposed progenitors

Tightly regulated release of ROS from healthy mitochondria plays a role in modulating signaling cascades, whereas dysfunctional mitochondria produce excess ROS that contribute to oxidative stress. Mitochondrial transmembrane polarization in undifferentiated progenitors was measured by the JC-1 and TMRE assays. A loss of membrane potential in BPS-exposed progenitors was apparent by a higher ratio of cationic JC-1 monomers relative to the J-aggregates formed when JC-1 enters the mitochondria (Figure 6A) and a lower accumulation of the cationic dye, TMRE (Figure 6B). Mitochondrial-derived superoxide (O_2_^−^) was increased in BPS-exposed progenitors (Figures 6C-E), while co-treatment of progenitors with the mitochondria-targeted antioxidant, MitoQ, markedly attenuated lipid droplet formation on day 4 of differentiation and abolished BPS-induced potentiation of adipogenesis (Figure 6F). Similarly, MitoQ attenuated the adipogenesis-enhancing effects of BPA and BPF (Supplementary Figure 2C &D). Co-treatment with the mitochondria-targeted antioxidant, SS31, which also protects against membrane depolarization, attenuated lipid droplet formation in differentiated progenitors exposed to BPS without having an impact on control cells (Figure 6G). Together, these data highlight mitochondria as sources of ROS that play a role in the regulation of adipogenesis and in the bisphenol-induced adipogenic response.

**Figure 6:**
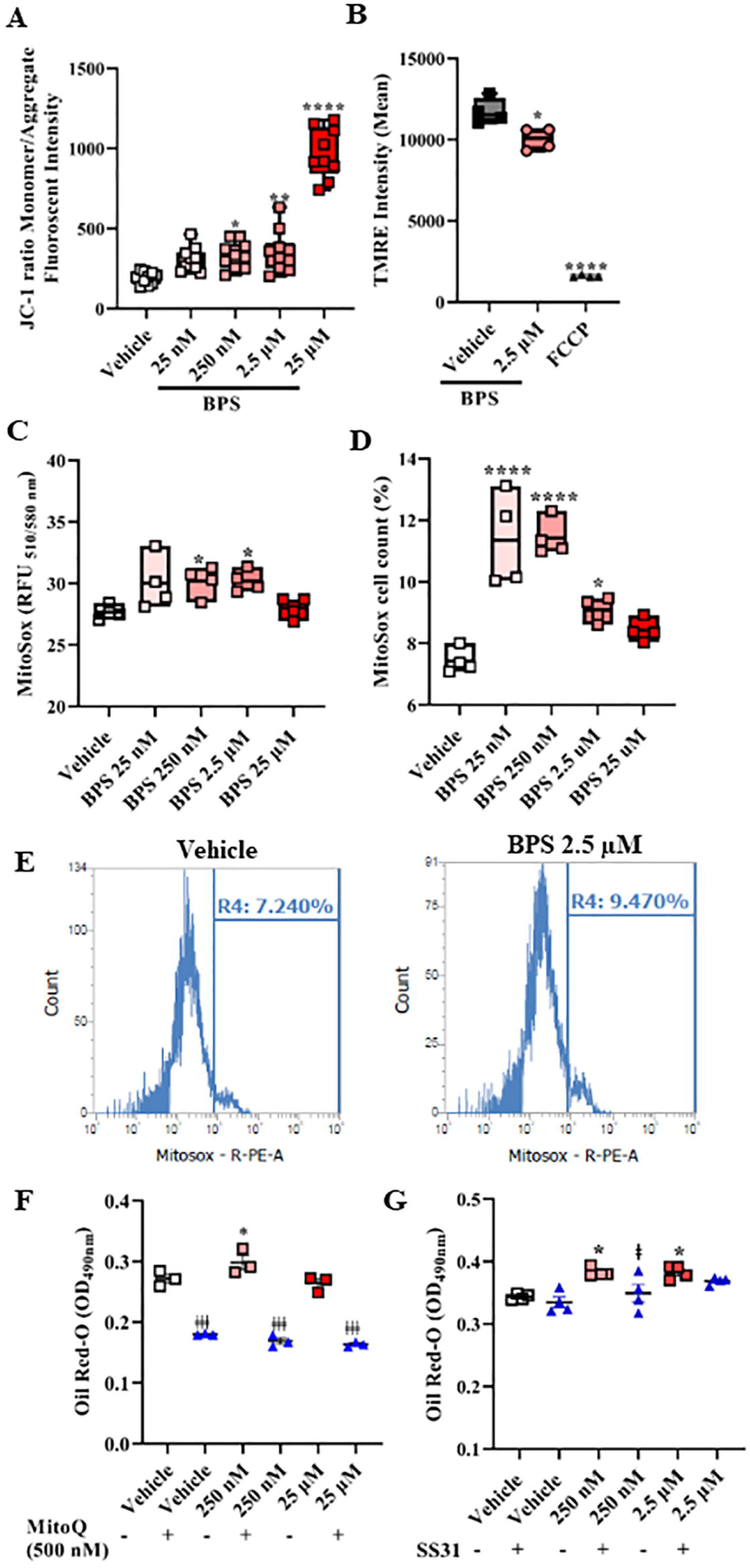
Mitochondria are important sources of ROS in BPS-exposed progenitors. Degree of depolarization calculated as the ratio of 5, 5’, 6, 6’-tetrachloro-1, 1’, 3, 3’-tetraethylbenzimidazolylcarbocyanine iodide (JC-1) monomer (green) to aggregate (red) fluorescent signal in undifferentiated progenitors exposed to BPS for 48 hrs (A). Flow cytometric quantification of tetramethyl rhodamine ethyl ester (TMRE) fluorescence indicating mitochondria membrane polarization in progenitors exposed to BPS or mitochondrial oxidative phosphorylation uncoupler (FCCP) that induces mitochondria membrane depolarization (B). Determination of mitochondria-derived superoxide (O_2_^−^) in undifferentiated stem cells exposed to BPS for 24 hrs by measuring relative fluorescence units (RFU) with a plate reader (C) or flow cytometric analysis (D). Representative histograms of progenitors stained positive for MitoSOX (E). Quantification of lipid droplet staining after differentiation of progenitors exposed to BPS or vehicle in the presence or absence of mitochondria-targeted antioxidants, MitoQ (F) or SS31 (G). Each experiment was performed in cells isolated from 2-3 animals and absorbance of triplicates from each well were averaged. Differences were assessed by One-way ANOVA with Dunnett’s multiple comparison test to compared BPS vs. vehicle (* p < 0.05) or One-way ANOVA with Tukey’s post hoc test to compare vehicle or BPS with inhibitor vs. without inhibitor (‡ p < 0.05 vs. without inhibitor).

### Prenatal BPS exposure programs higher adiposity in adulthood

To determine the impact of *in vivo* BPS exposure on adiposity, an environmentally relevant dose of BPS (2 μg/kg body weight) was administered via glass water bottles to C57BL/6J mice housed in an environment absent of plastic enrichment. Exposure occurred during the early life window of adipocyte commitment and differentiation (Gd0-Pd21) by exposing pregnant and lactating dams or in postnatal life after weaning (Figure 7A & B). Exposure to BPS during early life resulted in higher whole-body fat mass in adult male offspring, but not female offspring (Figure 7C). In contrast, exposure to BPS during postnatal life did not influence adiposity in adulthood in either male or female mice (Figure 7D).

**Figure 7:**
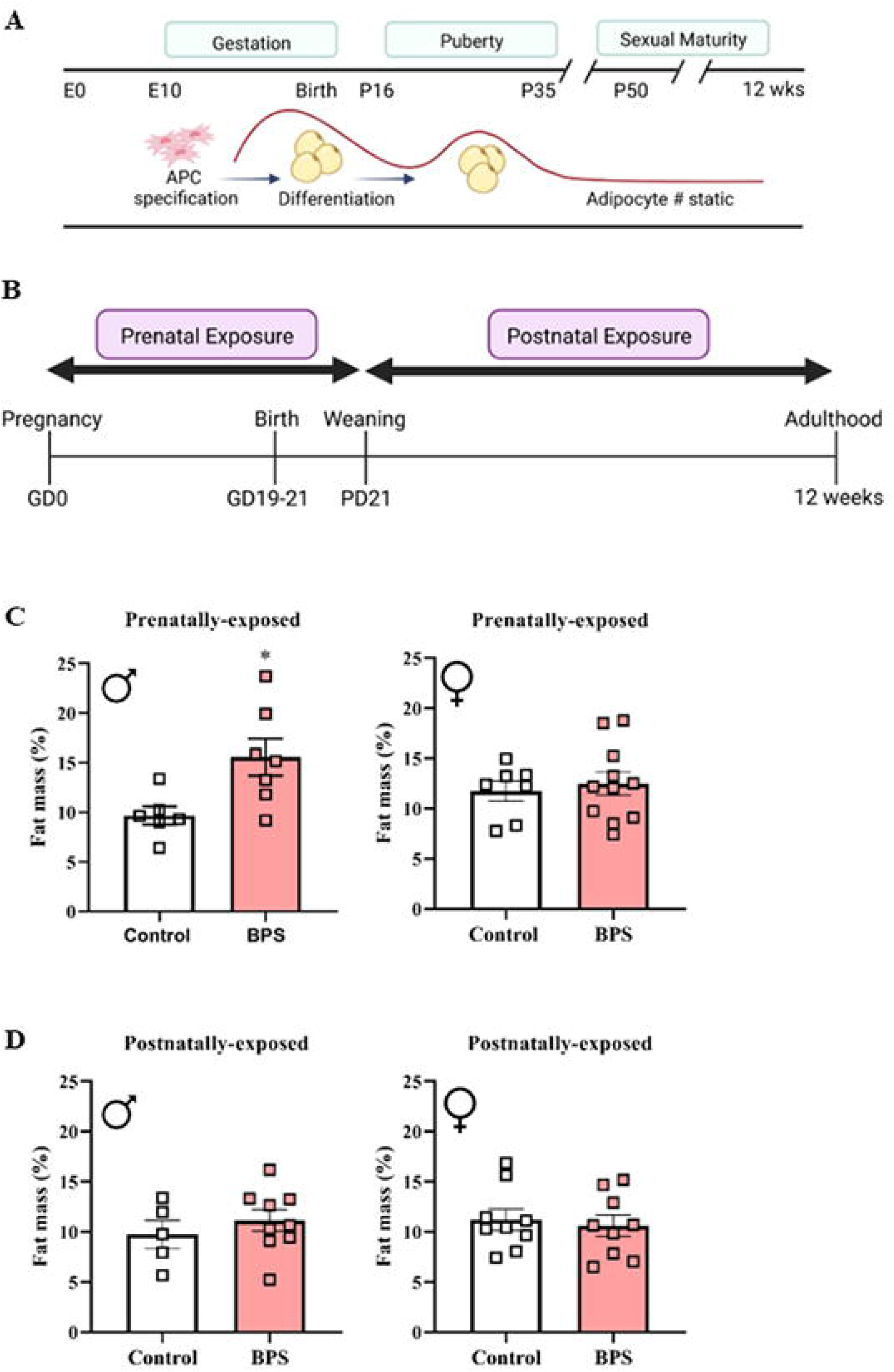
Intrauterine exposure to BPS programs an increase in later-life adiposity. In subcutaneous adipose depots, commitment of adipose progenitors cells (APC) to the adipocyte lineage occurs prior to birth, followed by rapid expansion of fat accumulation in the early postnatal period and during a second wave in puberty, with adipocytes numbers stabilizing thereafter (A). In C57BL6 mice, 2 μg/kg body weight BPS was administered to pregnant dams (prenatal exposure) starting on gestational day 0 (Gd0) and continuing until the pups weaned on postnatal day 21 (Pd21) or postnatal exposure to BPS was initiated on Pd21 (B). At 12 weeks of age prenatally exposed mice (C) or postnatally exposed mice (D) were assessed for whole-body fat mass content using TD-NMR. Fat mass of litter mates in prenatally-exposed mice is averaged for each sample (n = 7-11 litters). * p < 0.05 vs. control.

## DISCUSSION

Studies to-date have shown structural analogues marketed as safer substitutes for BPA to exhibit similar adipogenic properties as their predecessor (Ahmed and Atlas, 2016, Boucher et al., 2016). The adipogenesis-enhancing effects of BPA and its analogues have been attributed to their endocrine disrupting behaviour (Boucher et al., 2016), as they are classified as EDC that bind to estrogen, androgen, and glucocorticoid receptors, albeit with lower affinity and activity than endogenous ligands. Studies exploring endocrine pathways as mediators of the pro-adipogenic effects of bisphenols have yielded inconsistent findings with some reporting a role for estrogen and others showing a role for glucocorticoids (Ahmed and Atlas, 2016, Boucher et al., 2014). Results of the present study reveal a novel role for ROS in mediating the adipogenesis-promoting effect of BPS. Similar to BPS, BPA and BPF increased ROS production and had a potentiating effect on adipogenesis that was abolished by co-treatment with antioxidants. Therefore, cellular stress leading to heightened ROS production may be a common mechanism that explains the pro-adipogenic properties of BPA and its structural analogues. While it is well established that bisphenols and other EDC like phthalates promote oxidative stress in a variety of cell types (Špačková et al., 2020, Xie et al., 2020); to our knowledge this is the first study to explore the role of ROS in mediating the pro-adipogenic properties of EDC.

A handful of published studies demonstrate ROS to act as mediators of adipocyte differentiation (Schröder et al., 2009, Tormos et al., 2011); however, the role of redox signaling in adipogenesis has not received wide attention. Our data substantiate these studies as exposure of progenitors to low concentrations of O_2_^−^ generators enhanced differentiation, while antioxidant treatment inhibited adipogenesis in the absence of BPS. A non-monotonic response to increasing concentrations of BPS or ROS generators was observed, such that the increased differentiation at lower concentrations was attenuated at the highest dose. Cell viability was unaffected at BPS concentrations that enhanced adipogenesis, but reduced at the highest concentration of BPS, suggesting that the nonmonotonicity is due to cytotoxicity at higher levels of ROS production. Thus, cellular ROS promote adipocyte differentiation within a range of physiological concentrations below the cytotoxic threshold.

When produced at high levels in response to cellular stress, ROS damage macromolecules and contribute to cellular dysfunction, but at physiological levels, act as second messengers in signaling cascades that regulate a wide variety of cellular processes. ROS derive from several sources including mitochondria and enzymes such as the NADPH oxidases and xanthine oxidase. Silencing of NADPH oxidase 4 (Nox4) by RNA interference in bone marrow-derived mesenchymal stem cells or by siRNA in 3T3-L1 cells, abrogated lipid droplet formation after differentiation, implicating Nox4-derived hydrogen peroxide (H_2_O_2_) in the regulation of adipogenesis (Kanda et al., 2011, Schröder et al., 2009). Our findings cannot rule out a role for H_2_O_2_ as antioxidants and O_2_^−^ generators like DMNQ and menadione will influence levels of both O_2_^−^ and H_2_O_2_, as the former quickly dismutates to H_2_O_2_. Our data show that BPS exposure increases DCFDA staining of cellular ROS as well as O_2_^−^ levels inside mitochondria detected by MitoSox. Mitochondria are both generators and targets of ROS and crosstalk with enzymatic sources of ROS to promote an environment of oxidative stress. Thus, while our findings show mitochondria to be important contributors to BPS-induced oxidative stress, contributions from other sources of ROS cannot be ruled out.

Our data support studies showing that mitochondria-derived ROS are required for the induction of adipogenesis (Carrière et al., 2003, Tormos et al., 2011), as mitochondria-targeted antioxidants attenuated differentiation even in the absence of BPS. An increase in oxygen consumption during activation of the adipogenic program is driven by mitochondria biogenesis leading to an increase in mitochondria content (Li et al., 2017). Distinct subpopulations of mitochondria tethered to lipid droplets support formation and expansion of lipid droplets (Benador et al., 2019), the final steps in terminal differentiation. Mitochondria are major sources of ROS as complexes of the electron transport chain (ETC) leak electrons that combine with oxygen to produce O_2_^−^. The increase in oxidative phosphorylation that occurs during adipocyte differentiation is accompanied by an increase in ROS production (Tormos et al., 2011, Zhang et al., 2013). Decreasing mitochondria respiration by inhibiting the ETC abrogated differentiation of human mesenchymal stem cells into adipocytes (Zhang et al., 2013). Interestingly, a study by Tormos *et al*. showed that knockdown of a complex III subunit required for O_2_^−^ production attenuated differentiation of adipocyte progenitors, whereas adipogenesis was unaffected when complex III-derived O_2_^−^ was maintained with knockdown of a complex II subunit (Tormos et al., 2011). These findings demonstrate that an increase in mitochondria-derived ROS are not only by-products of differentiation but play a direct role in regulating adipogenesis that is independent of oxidative phosphorylation. Findings of the current study show that increased mitochondria-derived ROS in BPS exposed undifferentiated stem cells was accompanied by a loss of mitochondria membrane potential, indicative of mitochondria injury. Therefore, disruption of mitochondria leading to moderate levels of ROS may potentiate differentiation, while more extensive mitochondrial damage and oxidative stress may interfere with the mitochondrial dynamics required for successful differentiation.

Moderate mitochondria membrane depolarization has been shown to serve as a protective mechanism that preserves energetic capacity while mitigating mitochondria-derived ROS under conditions of ischemia (Vyssokikh et al., 2020). Therefore, it is possible that the mitochondria membrane depolarization in BPS-exposed progenitors is a compensatory response to cellular stress.

Although *in vitro* studies consistently show an adipogenesis-promoting effect of BPA and its analogues, human and animal studies have yielded inconsistent results with respect to the relationship between bisphenol exposure and adiposity (Callaghan et al., 2020). Adipogenic responses *in vitro* do not necessarily reflect adipogenesis *in vivo*, the latter influenced heavily by the microenvironment of the adipose depot. Further, exposure to bisphenols have pleiotropic effects with the potential to impact metabolism and adiposity through multiple mechanisms. Nevertheless, several epidemiological studies have demonstrated relationships between urinary levels of BPA or its analogues and components of the metabolic syndrome including abdominal obesity, hypertension, and insulin resistance (Callaghan et al., 2020). Most human studies use single timepoint measurements of bisphenol levels and are unable to assess low dose affects due to ubiquitous exposure. Further, humans are not exposed to single toxicants, but rather toxicant mixtures, and correlates to exposure such as consumption of processed foods are confounding variables. Animal studies vary widely in the dose, timing, and duration of bisphenol exposure, some reporting prenatal BPA exposure to result in increased adiposity and some reporting fetal growth restriction (Rubin et al., 2019). The current study demonstrated that BPS exposure in pregnant and lactating dams leads to an increase in adiposity in adult male offspring, while postnatal exposure had no impact on fat mass. Seminal studies by Graff and others show that the progenitor pool giving rise to adipocytes during postnatal expansion of subcutaneous adipose depots are specified to the adipocyte fate before birth (Jeffery et al., 2015, Jiang et al., 2014, Wang et al., 2013, Wang et al., 2015). Thus, our data show that a programmed predisposition to high adiposity occurs when exposure coincides with this critical window of adipogenesis. However, it is also possible that higher adiposity in adulthood is a secondary effect of other developmental perturbations. For instance, some studies report maternal bisphenol exposure to result in fetal growth restriction, which is well known to cause rapid catch-up growth and program a predisposition to obesity. The sexual dimorphism in programming of adiposity by early life BPS exposure suggests an interaction of BPS with endocrine pathways that warrants further investigation.

The dose used for *in vivo* exposure was below the tolerable daily intake (TDI) for BPA (4μg/kg body weight/day) established by the European Food Safety Authority (EFSA). Currently, there are no established limits for BPA analogues. Recent findings from China show that ingestion of BPS from contaminated food, the primary route of human exposure, ranged between 5.74 to 56.9 ng/kg body weight/day, exceeding that of BPA intake (Zhang et al., 2022). In Alberta, Canada, daily 24 hr intake of BPS in pregnant women reached up to 14 nM/kg body weight, approaching the TDI of BPA (Liu et al., 2018). We acknowledge that prenatally-exposed and postnatally-exposed adult mice were not exposed to the same level of BPS; however, our approach reflects the real-world scenario where the fetus and lactating infant is exposed to a fraction of the mother’s intake of BPS. Nevertheless, our findings suggest that BPA analogues are obesogenic and reveal increased ROS production as a contributing mechanism.

## INNOVATION

Pro-adipogenic effects of BPA are well characterized; however, little is known regarding new structural analogues that are replacing BPA since introduction of new regulatory restrictions and heightened awareness of its adverse health effects. Molecular pathways underlying the adipogenesis-promoting properties of bisphenols remain unresolved, and studies to-date have almost exclusively focused on their endocrine disrupting properties. Herein, we undertook extensive investigation that substantiates existing data identifying ROS act as signaling molecules in the regulation of adipocyte differentiation and highlight ROS as a unifying mechanism that explains the pro-adipogenic properties of BPA and its analogues that is independent of endocrine disruption (Fig. 1).

## MATERIAL AND METHODS

### Isolation and differentiation of adipocyte progenitor cells

All animal procedures were approved by the University of Calgary Animal Care Committee (AC19-0006) and conducted in accordance with guidelines by the Canadian Council on Animal Care Ethics. At 6-8 weeks of age, C57BL/6J male mice were sacrificed for harvesting of inguinal SAT (iSAT), which was digested in a 1mg/ml Collagenase I (Worthington Biochemical, Lakewood, New Jersey, cat no: LS004194) digestion buffer in HBSS with 100 mM HEPES and 1.5% BSA. The pelleted stromal vascular fraction (SVF) was treated with RBC lysis buffer (Alfa Aesar, Tewksbury, Massachusetts, cat no: J62150) resuspended in preadipocyte growth medium (Cell Applications, San Diego, California, cat no: 811), and cultured at 37 °C and 5% CO_2_. After a single passage, cells were seeded, and differentiation was induced 48 hours after contact inhibition (Day 0) until day 5 when differentiation medium (Cell Applications, cat no: 811D) was replaced with maintenance medium (Cell Applications, cat no: 811M). Prior to contact inhibition (90% confluency) cells were treated with various doses of BPS (Sigma-Aldrich, Oakville, Ontario, cat no: 103039), BPA (Sigma-Aldrich, cat no: 239658), BPF (Sigma-Aldrich, cat no: 51453) or ROS generators or vehicle (DMSO), in the presence or absence of antioxidants. Menadione (Sigma-Aldrich, cat no: M5625) and 2,3-dimethoxy-1,4-napthalenedione (DMNQ, Sigma-Aldrich, cat no: D5439) were used as ROS generators. Lipid droplets were stained with an Oil Red O solution (Sigma-Aldrich, cat no: O0625) or 2 μM BODIPY (Thermo Fisher Scientific, Mississauga, Ontario, cat no: D3922), captured using a Nikon Eclipse Ts2 microscope and processed with NIS-Elements D 5.11.00. After thorough washing, Oil-Red O-stained cells were incubated with a lysis buffer and the absorbance of the eluted dye quantified in triplicates at 490nm on a spectrophotometer (Biotek 800TS).

### Prenatal and postnatal exposure to BPS

After weaning, female C57BL/6J mice (Jackson Laboratories,, Bar Harbor, Maine, stock #: 000664) were transferred to cages equipped with glass water bottles (Lab Products LLC, Seaford, Delaware, cat no: 30800) and absent of plastic enrichment to eliminate background exposure to plasticizers. For prenatal exposure, 12-week-old dams were randomized to BPS drinking water (2 μg/kg body weight/day) or vehicle control, started on a phytoestrogen-low diet (Envigo, Indianapolis, Indiana, Teklad diet: 2020) and mated with males. Exposure was initiated at gestational day 0 (Gd0) after confirmation of pregnancy by observation of a copulatory plug. Dams were allowed to deliver spontaneously, and litters culled to a maximum of 6 to minimize litter effects. Exposure continued through lactation until postnatal day 21 (Pd21), at which time pups were weaned and maintained on a plastic-reduced environment. For postnatal exposure, littermates were randomized to BPS or vehicle drinking water after weaning. At 12-weeks of age, mice were sacrificed and whole-body fat mass measured with a TD-NMR body composition analyzer (Bruker, Billerica, Massachusetts, LF90II). Fat mass of litter mates was averaged for prenatally-exposed offspring to control for litter effects, while litter mates were compared in postnatally-exposed mice.

### RNAseq and Pathway Analysis

RNA samples were assessed with TapeStation and Qubit assays and 1000ng RNA was used in library preparation using the NEBNext Ultra II directional RNA library prep kit for Illumina, along with the NEBNext Poly(A) mRNA Magnetic Isolation Module. Final libraries were assessed with the Kapa library quantification qPCR assay and sequencing performed on the NextSeq 75 cycle high output run. The sequencing run passed all QC metrics. Samples were quantified using Kallisto v0.42.4 against the NCBI RefSeq transcriptome (GRCm38) (Bray et al., 2016). A linear regression model was used to determine differentially expressed transcripts, with terms for treatment as well as sample admixture proxy markers identified in principal component analysis. Differentially expressed genes (DEG) were considered as those with a false discover rate (Benjamini-Hochberg corrected p-value) < 0.05 under the Wald test in Sleuth v0.30.0 (Pimentel et al., 2017). DEGs were subsequently annotated and analyzed using the Molecular Signatures Database (MSigDB) hallmark pathway analysis using Enrichr (https://maayanlab.cloud/Enrichr/), a comprehensive gene set enrichment analysis web server. A heatmap of DEGs involved in adipogenesis and the Nrf2-mediated oxidative stress response was created using R studio software (2022.02.0+443). The potential interaction between DEGs involved in these two pathways was presented using Interaction Network Analysis using the Path Explorer Tool of Qiagen IPA software. These data have been deposited in NCBI’s Gene Expression Omnibus and are accessible through GEO Series accession number GSE213781 (https://www.ncbi.nlm.nih.gov/geo/info/linking.html).

### Quantitative real time PCR

Total RNA was isolated using the RNeasy Mini Kit (Qiagen, Hilden, Germany, cat no: 74004), assessed for integrity using a TapeStation RNA assay (Agilent, Santa Clara, California) and quantified using a N50 Nanophotometer (Implen Inc., Westlake Village, California). Following DNase treatment, cDNA synthesis was performed using a High-Capacity cDNA Reverse Transcription kit (Applied Biosystems, Walthan, Massachusetts, cat no: 4368814). cDNA products were run in triplicates for Quantitative real time PCR using a QuantStudio 5 Real-time PCR System (Applied Biosystems, cat no: 4368814) and Powerup SYBR green master mix (Applied Biosystems, cat no: A25741). Primers were designed using the NCBI/Primer Blast tool (Supplementary Table 1). Target mRNA expression was normalized to β-actin and fold change calculated using 2^−ΔΔCt^ method.

### Western blot

Proteins were extracted from cell lysates in buffer containing a protease and phosphatase inhibitor cocktail (Thermo Fisher Scientific, cat no: 78440) and resolved in precasted NuPAGE 4%–12% Bis-Tris polyacrylamide gels (Invitrogen, Oregon, cat no: NPO321) and then transferred to Amersham Hybond PVDF membranes (GE Healthcare Life Sciences, Illinois, cat no: 10600069) at 100V for 2 hr. The membrane was then washed, blocked with 5% blotting grade blocker (Bio Rad, Hercules, California, cat no: 1706404) for 1 hr and probed with primary antibodies [Cell Signaling, Danvers, Massachusetts, ADIPOQ (2789S); C/EBPβ (3087S); SCD1 (2794S); FAPB4 (2120S); GAPDH (2118S)]. After washing and re-probing with a horseradish peroxidase-conjugated secondary antibody (Thermo Fisher Scientific, cat no: A11077), bands were visualized with the iBright CL1500 Imaging system (Applied Biosystems).

### ROS detection

For the detection of cellular ROS, cells were prestained with 2′,7′-dichlorodihydrofluorescein diacetate (DCFDA, Sigma-Aldrich, cat no: D6883), washed and incubated in BPS at 37 °C for 45 min. CellROX™ deep red dye (5μM, 640/665 nm, Invitrogen, cat no: C10422) and MitoSOX™ Red dye (2.5 μM, 510/580 nm, Invitrogen, cat no: M36008) were used to detect cellular ROS and mitochondria-derived ROS, respectively, after 24 or 48 hr of BPS exposure. Relative Fluorescent Unit (RFU) was measured with a spectrophotometer (SpectraMax M2) and normalized to background fluorescence and protein concentration. Flowcytometric analysis was carried out using an Attune® Acoustic Focusing Cytometer (ThermoFisher Scientific) after cells were dissociated using 5 mM EDTA in HBSS, centrifuged at 500g and resuspended in HBSS. Stained cells were imaged with the 60X objective using a Nikon Ti Eclipse Widefield microscope, employing the DAPI, CY5, and CY4 filters.

### Mitochondria function

After 24 hrs of BPS exposure, cells were incubated in 2 μM JC-1 (5, 5’, 6, 6’-tetrachloro-1, 1’, 3, 3’-tetraethylbenzimidazolylcarbocyanine iodide, Invitrogen, cat no: T3168) at 37°C for 15 min. After washing, fluorescence signals were measured at 514/529 (green, monomer) and 514/590 (red, aggregate) using SpectraMax^®^ M2 and the mitochondria membrane potential expressed as the ratio of green to red fluorescence. The mitochondria membrane potential was also assessed in live cells exposed to BPS for 24 hrs using tetramethyl rhodamine ethyl ester (TMRE, Thermo Fisher Scientific, cat no: T669) staining. Live cells were collected in HBSS containing 5 mM EDTA, washed and resuspended in media and the TMRE mean fluorescent signal quantified on a flow cytometer.

### Statistical Analysis

Statistical analyses were performed using GraphPad Prism 9. Two-tailed student’s t-test or One-way ANOVA with Dunnett’s or Tukey’s multiple comparison test were used to evaluate differences between groups. All data in this study are represented as the mean ± standard error of mean (SEM) or maximum, minimum, and median. P values <0.05 are considered statistically significant. An electronic laboratory notebook was not used.

## LIST OF ABBREVIATIONS

(ADIPOQ): Adiponectin
(Fos): AP-1 transcription factor subunit
(BPA): Bisphenol A
(BPF): bisphenol F
(BPS): bisphenol S
(Cat): catalase
(C/EBPβ): CCAAT/enhancer binding protein beta
(EDC): endocrine disrupting chemical
(ETC): electron transport chain
(FABP4): fatty acid binding protein 4
(FASN): fatty acid synthase
(GLUT4): glucose transporter type 4
(Gpx1): Glutathione peroxidase 1
(Klf4): Kruppel-like factor 4
(Nr6a1): nuclear receptor sub-family 6 group A member 1
(PPARγ): peroxisome proliferator-activated receptor gamma
(Ptgs2): prostaglandin endoperoxidase synthase 2
(ROS): reactive oxygen species
(RBC): red blood cell
(SAT): subcutaneous adipose tissue
(SCD1): stearoyl-CoA desaturase 1
(SVF): stromal vascular fraction
(SOD): superoxide dismutase

## FIGURE LEGENDS

**Supplementary Figure 1: Cell viability is not compromised over the range of BPS concentrations that promote adipogenesis**. Quantification of MTT, as an indicator of cell viability, in undifferentiated progenitors exposed to various concentrations of BPS for 24 hrs (A) or 48 hrs (B). Differences compared by One-way ANOVA with Dunnett’s post hoc test. * p < 0.05 BPS vs. vehicle.

**Supplementary Figure 2: Enhanced lipid droplet formation in stem cells exposed to BPA or BPF is attenuated with a mitochondria-specific antioxidant**. Quantification of eluted Oil-Red O dye in differentiated progenitors after exposure to various concentrations of BPA (A) or BPF (B). Lipid droplet formation was measured in presence or absence of MitoQ to assess the role of mitochondria-derived ROS in mediating the effect of BPA (C) or BPF (D) on adipogenesis. Differences were assessed by One-way ANOVA with Dunnett’s multiple comparison test to compared BPS vs. vehicle (* p < 0.05) or One-way ANOVA with Tukey’s post hoc test to compare vehicle or BPS with inhibitor vs. without inhibitor (‡ p < 0.05 vs. without inhibitor).

## ACKNOWLEDGEMENTS

We would like to thank Paul Gordan and the Centre for Health Genomics and Informatics at the University of Calgary for support in analyzing the RNAseq data. We also thank Dr. Steven Greenway from the Libin Cardiovascular Institute for providing the mitochondria-targeted antioxidant, SS31.

## AUTHOR CONTRIBUTIONS

RDS contributed to the generation of hypotheses, experimental design, data collection, analysis, and manuscript writing. JW, MC, TS, LC, SE and SAH contributed to data collection and analysis. AH, VP and TS provided support for mitochondria analyses, while PC and LS provided support for imaging.

