## Supplementary figures and images for "Potentiation of Adipogenesis by Reactive Oxygen Species is a Unifying Mechanism in the Pro-adipogenic Properties of Bisphenol A and its New Structural Analogues"

### Supplementary Figure 1

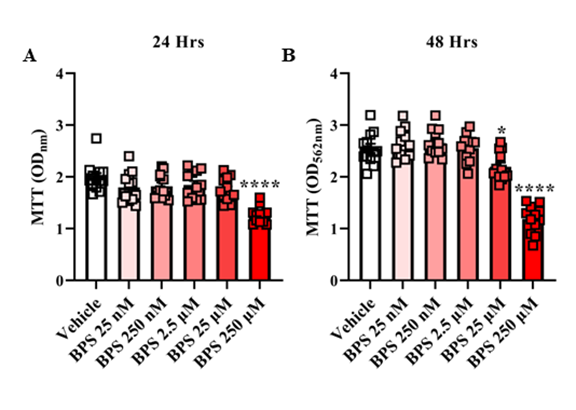

### Supplementary Figure 2

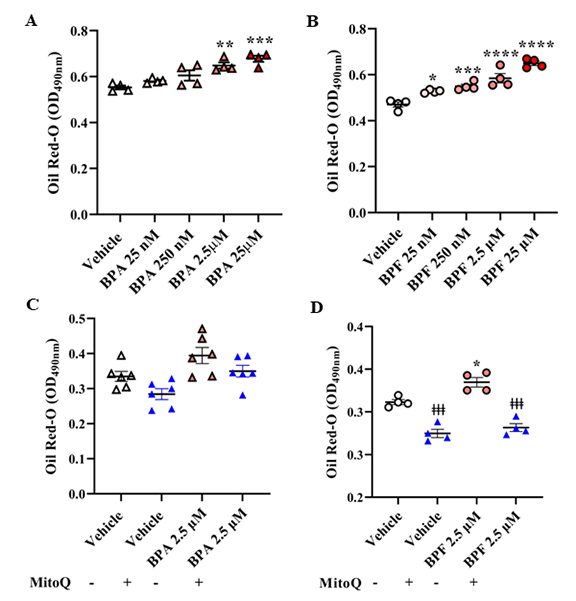
