## Supplementary Table 1 for "Potentiation of Adipogenesis by Reactive Oxygen Species is a Unifying Mechanism in the Pro-adipogenic Properties of Bisphenol A and its New Structural Analogues"

**Supplementary Table 1:** Primer pairs

| **Gene** | | **Sequence** | **Accession** |
| --- | --- | --- | --- |
| *Klf4* | Forward | CATTAATGAGGCAGCCACCTG | NM_010637.3 |
|  | Reverse | GCTTCATGTGAGAGAGTTCCT |  |
| *Fos* | Forward | CTTCGACCATGATGTTCTCG | \| NM_010234.3 |
|  | Reverse | CTGGGGAATGGTAGTAGGAA |  |
| *Cat* | Forward | CTTTGACAGAGAGCGGATTC | NM_009804.2 |
|  | Reverse | CTTTGCCTTGGAGTATCTGG |  |
| *Sod3* | Forward | ACCCAGAAATCTTTTCACGC | NM_011435.3 |
|  | Reverse | AGCAGACTCAAAGACTAGGG |  |
| *Gpx1* | Forward | AGTCCACCGTGTATGCCTTC | NM_008160.6 |
|  | Reverse | CAGATCGTTCATCTCGGTGT |  |
| *Nr6α1* | Forward | ATCATCCAGTAGGTCTGTGG | AF390896.1 |
|  | Reverse | GTCACAGCATACCCATCTTC |  |
| *β-actin* | Forward | GATCAAGATCATTGCTCCTCCT | NM_007393.5 |
|  | Reverse | GTAACAGTCCGCCTAGAAGC |  |
| *C/ebpβ* | Forward | TGATGCAATCCGGATCAAAC | NM_001287738.1 |
|  | Reverse | CCGCAGGAACATCTTTAAGT |  |
| *Pparγ* | Forward | GCTGTTATGGGTGAAACTCT | EF062476.1 |
|  | Reverse | TGATGTCAAAGGAATGCGAG |  |
| *Glut4* | Forward | GGTGGCATGATCTCTTCCTT | NM_001359114.1 |
|  | Reverse | CCTGATGTTAGCCCTGAGTA |  |
| *Scd1* | Forward | CAAGAGTAGCTGAGCTTTGG | NM_009127.4 |
|  | Reverse | AATGCATCATTAACACCCCG |  |
| *Fasn* | Forward | AGGGTACAAAGGTGTTCAGA | NM_007988.3 |
|  | Reverse | CTCATGACGAGTGCTGTAAG |  |
| *Ptgs2* | Forward | AAAACCGTGGGGAATGTATG | [M64291.1](https://www.ncbi.nlm.nih.gov/nucleotide/M64291.1?report=genbank&log$=nucltop&blast_rank=16&RID=KBK76P5H013) |
|  | Reverse | GGGCTTCAGCAGTAATTTGA |  |

**Footnotes:** AP-1 transcription factor subunit (Fos); catalase (Cat); CCAAT/enhancer binding protein beta (C/EBPβ); fatty acid binding protein 4 (FABP4); fatty acid synthase (FASN); glucose transporter type 4 (GLUT4); Glutathione peroxidase 1 (Gpx1); Kruppel-like factor 4 (Klf4); nuclear receptor sub-family 6 group A member 1 (Nr6a1); peroxisome proliferator-activated receptor gamma (PPARγ); prostaglandin endoperoxidase synthase 2 (Ptgs2); stearoyl-CoA desaturase 1 (SCD1); superoxide dismutase (SOD).
